## Supporting Information for "Single-Molecule Imaging and Microfluidic Platform Reveal Molecular Mechanisms of Leukemic Cell Rolling"

**Contents**

Supporting Methods

Figures S1 to S26

Supporting Notes 1 to 7

Captions for Movie S1 to S7

Supporting References

### Supporting Methods

**Fragmentation and fluorescence labeling of the antibodies.** For the digestion and purification of the anti-PSGL-1 antibody (KPL-1 clone, IgG1), we used Pierce Mouse IgG1 Fab and F(ab′)2 Preparation Kit or Pierce Mouse IgG1 Fab and F(ab′)2 Micro Preparation Kit (Thermo Scientific). The kit uses immobilized ficin protease to efficiently digest mouse IgG1 into Fab or F(ab')2 fragments, depending on the concentration of cysteine and solution pH. Briefly, the digestion buffer was prepared by dissolving 43.9 mg of cysteine•HCl in 10 ml of the supplied Mouse IgG1 digestion buffer (pH = 5.6) to produce Fab fragments. Then, the immobilized enzyme was dispensed into the spin column, centrifuged and washed using the prepared digestion buffer. The antibody (1 mg $ml^{-1}$, 125 µl) was desalted using the accompanied Zeba column. The flow through of at least 100 µl that contained the antibody was incubated with the immobilized enzyme for 3-5 hours at 37$^{o}$C on an end-over-end mixer, which digested the antibody into Fab fragments. After the incubation, the column was put into a 2 ml collection tube and centrifuged at 5000 × g for 1 min to collect the digested antibodies. The collected flow through was incubated with protein A column for 10 min at room temperature on an end-over-end mixer, followed by centrifugation at 1000 × g for 1 min to separate the Fab fragments from Fc and undigested antibodies. The flow through that contained Fab fragments was collected. After the fragmentation and purification, concentrations of the anti-PSGL-1 antibody (KPL-1 clone, IgG1) and its fragments were determined by measuring the absorbance at 280 nm using UV-vis spectrophotometer (Thermo Scientific, NanoDrop 2000).

Fluorescence labeling of the Fab fragment of the anti-PSGL-1 antibody by the Alexa-Fluor dyes conjugated to N-hydroxysuccinimide (NHS) was conducted in a manner similar to the labeling of the whole antibody. The Fab fragment and the dye were mixed at the mixing ratio of thirty to one, which corresponds to the dye to antibody molar ratio of approximately 120 to 1. The degree of labeling with this mixing ratio was in the range of 1 – 3 dyes per Fab fragment.

**Flow cytometry.** Binding specificity of the mouse anti-human PSGL-1 and mouse anti-human CD44 antibodies, which were used for the fluorescence imaging experiments, to PSGL-1 and CD44 on KG1a cells were evaluated by flow cytometry. KG1a cells ($10^6$ cells $ml^{-1}$) were incubated with either anti-human PSGL-1 antibody (clone KPL-1, Ms IgG1, $_\kappa$), anti-human CD44 antibody (clone 515 Ms IgG1, $_\kappa$, or 2C5 Ms IgG2a) or purified mouse IgG1κ isotype (BioLegend) at a final concentration of 10 μg $ml^{-1}$ in HBSS at 4°C for 30 min. Subsequently, the KG1a cells were incubated with Alexa-Fluor-488–conjugated goat anti-mouse antibody (5 μg $ml^{-1}$, IgG, Invitrogen) in HBSS at 4°C for 20 min. For the secondary antibody control, KG1a cells ($10^6$ cells $ml^{-1}$) were incubated with Alexa-Fluor-488–conjugated goat anti-mouse antibody (5 μg $ml^{-1}$) in HBSS at 4°C for 20 min. The fluorescence intensity was determined using a FACSCanto flow cytometer (Beckman Dickenson).

**Western Blot Analysis.** WT KG1a, scrambled, or CD34 siRNA knockdown cells were lysed using a cell lysis buffer containing 88% NP40 (Invitrogen™ Novex™, Fisher Scientific), 10% protease inhibitor (Pierce™, Thermo Scientific), 1% PMSF, and 1% Phosphatase inhibitor (Halt™, Thermo Scientific) at 4°C for 1 h. The whole cell lysate was collected and incubated with NuPAGE LDS sample buffer (Invitrogen) and 10% β-mercaptoethanol at 70°C for 10 min. The samples were then run on an SDS-PAGE gel prior to being transferred to a PVDF membrane. The PVDF membrane was blocked overnight at 4°C using Tris-buffered saline with Tween-20 (Cell Signaling Technology) containing 5% non-fat skim milk powder. The membrane was then washed and incubated with a mouse anti-human CD34 antibody (QBEND/10, BIO RAD). The membrane was then washed and immunoblotted with HRP-conjugated secondary antibodies prior to imaging.

**Supporting Figures**

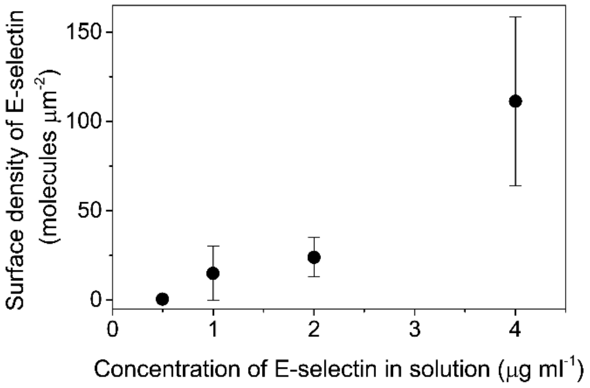


**Figure S1.** Surface density of rh E-selectin molecules. The surface densities of the rh E-selectin molecules were determined after incubating the microfluidic chambers with HBSS buffer containing varied concentrations of the recombinant E-selectin at 4 °C overnight. The error bars show the standard deviations determined by at least four separate experiments.

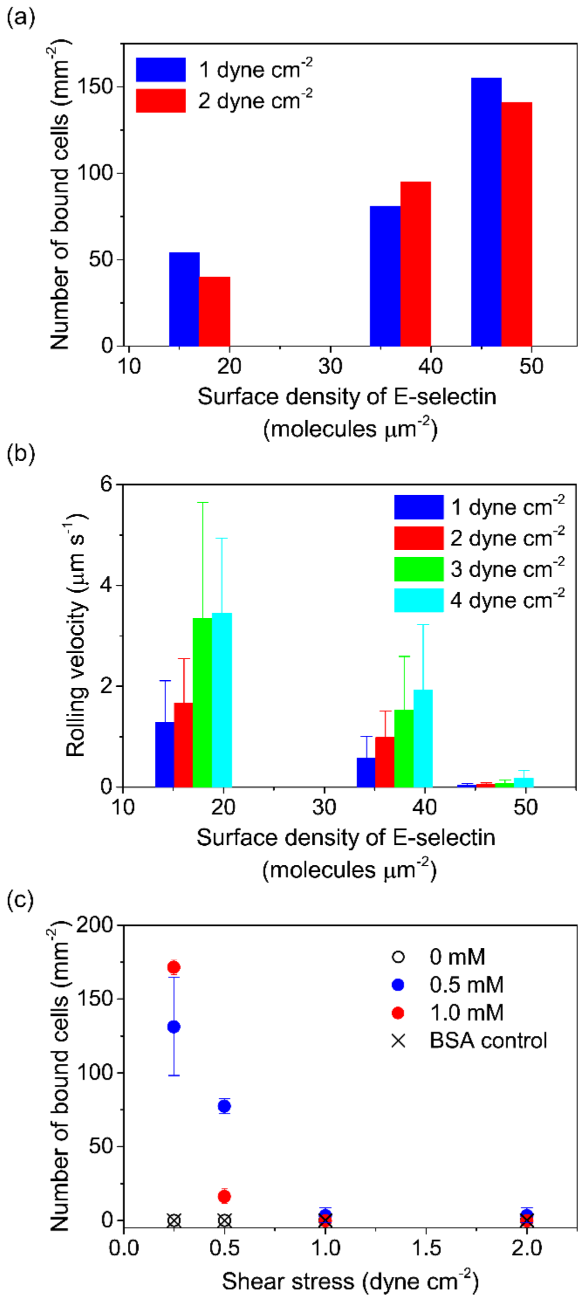


**Figure S2.** Characterization of the rolling behavior of KG1a cells on E-selectin. (a) Number of bound cells to the E-selectin surface at varied surface densities of the deposited rh E-selectin. The cells were injected into the microfluidic chambers at the shear stresses of either 1 or 2 dyne cm$^{-2}$ (0.1 or 0.2 Pa). (b) Rolling velocity of the cells on the E-selectin surface at different surface densities of the deposited rh E-selectin. The cells were injected into the microfluidic chambers at the shear stresses of either 1, 2, 3, or 4 dyne $cm^{-2}$ (0.1, 0.2, 0.3, or 0.4 Pa). The error bars show the standard deviations determined by at least six separate experiments. (c) Number of bound cells to the E-selectin surface at different applied shear stresses. The cells were injected into the microfluidic chambers at the calcium ion concentrations of either 0, 0.5, or 1.0 mM. BSA control (i.e. no E-selectin on the surface) is included as a negative control. The error bars show the standard deviations determined by at least three separate experiments.

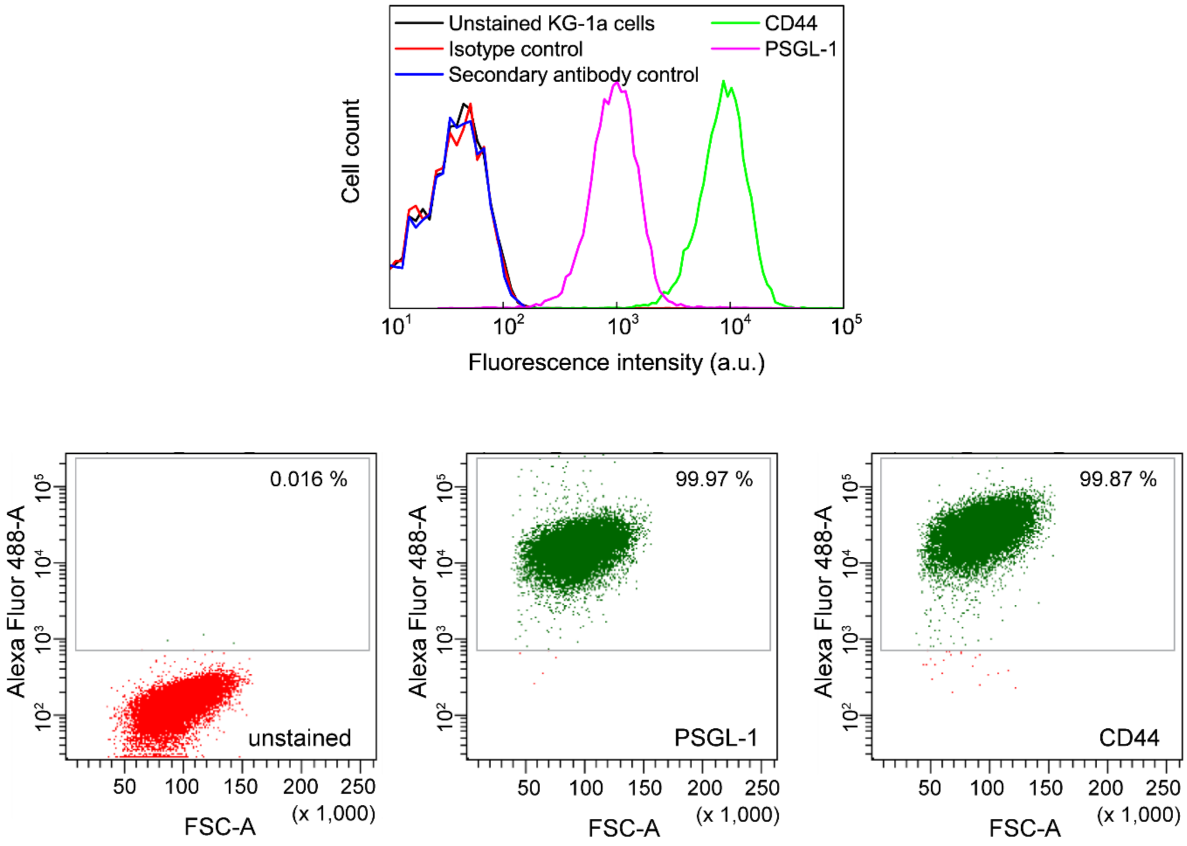


**Figure S3.** Flow cytometric analysis of the binding specificity of CD44 and PSGL-1 antibodies. CD44 (green) and PSGL-1 (magenta) on KG1a cells were immunostained by antibodies against CD44 and PSGL-1 (clone 515 for CD44 and clone KPL-1 for PSGL-1) and Alexa-Fluor-488-conjugated secondary antibody against the isotype of the CD44 and PSGL-1 antibodies (goat anti-mouse IgG). Unstained KG1a cells (black), isotype control labelled cells (red), and secondary alone-labelled cells (blue) were included as controls. This is a representative experiment of n=2 independent experiments.

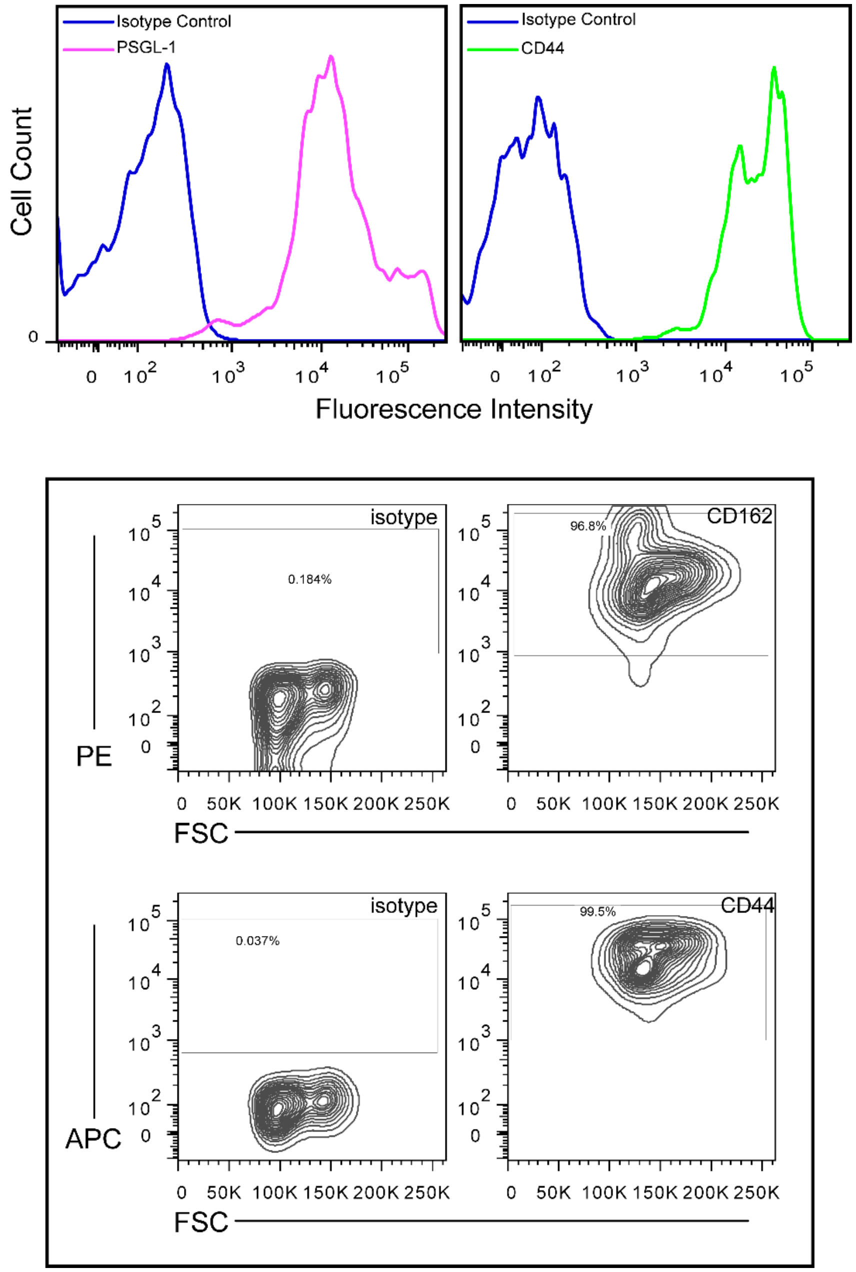


**Figure S4.** Assessment of E-selectin ligand expression on human CD34$^{pos}$-HSPCs. Primary human CD34$^{pos}$-HSPCs were stained for CD44 (green) and PSGL-1 (magenta) using antibodies specific to these antigens (anti-human CD44, clone 2C5 and anti-human PSGL-1, clone KPL-1) and analyzed by flow cytometry (FACS Canto and FlowJo). This is a representative experiment of n=2 independent experiments.

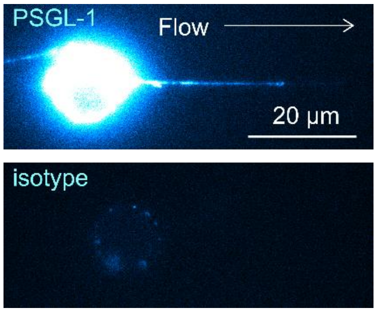


**Figure S5.** Binding specificity of the antibody characterized by fluorescence imaging. (top) Fluorescence image of PSGL-1 molecules on a KG1a cell that were immunostained by Alexa-Fluor-555-conjugated antibody (anti-PSGL-1 antibody, clone KPL-1). The labeled cells were injected into the rh E-selectin-deposited microfluidic chambers. The cells were injected into the chambers at a shear stress of 2 dyne $cm^{-2}$ (0.2 Pa). (bottom) Fluorescence image of an isotype control labelled KG1a cell that was injected into the fluidic chambers at conditions identical to those for the PSGL-1 immunostained cells.

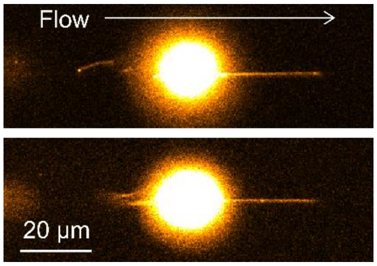


**Figure S6.** Formation of tethers and slings on KG1a cells while rolling over E-selectin. Examples of the fluorescence images of the cell membrane (stained by Vybrant DiO dye) captured during cell rolling over the surface-deposited rh E-selectin molecules. The cells were injected into the chambers at a shear stress of 2 dyne $cm^{-2}$ (0.2 Pa).

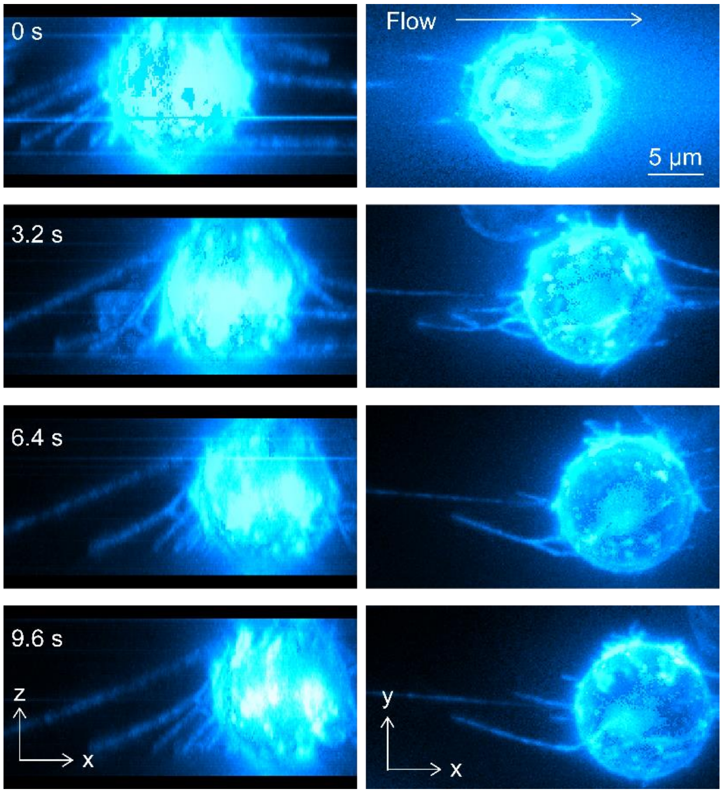


**Figure S7.** 3D views of the tethers and slings formed on a KG1a cell rolling over E-selectin by CD44. Side view (left) and top view (right) of the 3D reconstructed time-lapse fluorescence images of CD44 (immunostained by Alexa-Fluor-647-conjugated anti-CD44 antibody, clone 515) captured during cell rolling over the surface-deposited rh E-selectin molecules. The 3D images were reconstructed by recording fluorescence images of the cell at 53 different Z-axis positions with 0.5 μm step size. The cells were injected into the chambers at a shear stress of 2 dyne $cm^{-2}$ (0.2 Pa).

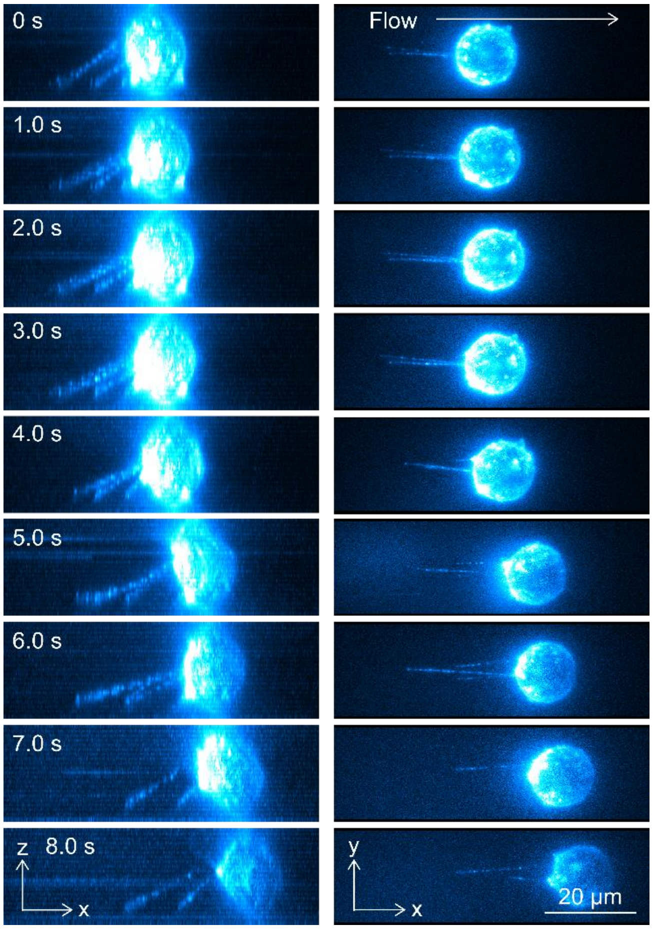


**Figure S8.** 3D views of the tethers formed on a KG1a cell rolling over E-selectin by PSGL-1. Side view (left) and top view (right) of the 3D reconstructed time-lapse fluorescence images of PSGL-1 (immunostained by Alexa-Fluor-555-conjugated anti-PSGL-1 antibody, clone KPL-1) captured during cell rolling over surface-deposited rh E-selectin molecules. The 3D images were reconstructed by recording fluorescence images of the cell at 33 different Z-axis positions with 1.0 μm step size. The cells were injected into the chambers at a shear stress of 2 dyne $cm^{-2}$ (0.2 Pa).

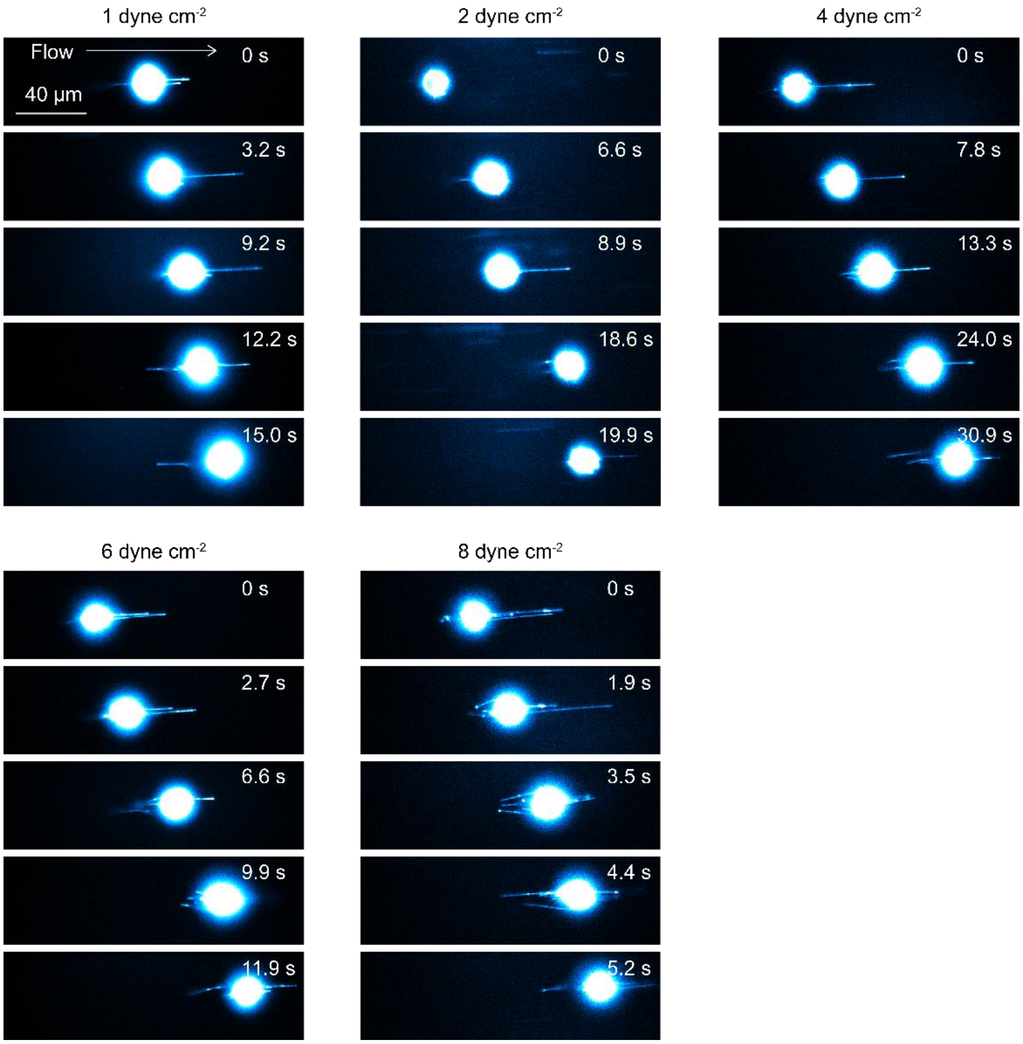


**Figure S9.** Shear force-dependent formation of the tethers and slings on KG1a cells rolling over E-selectin. Time-lapse fluorescence images of CD44 (immunostained by Alexa-Fluor-647-conjugated anti-CD44 antibody clone 515) captured during cell rolling over the surface-deposited rh E-selectin molecules. The cells were injected into the chambers at a shear stress of 1, 2, 4, or 8 dyne cm$^{-2}$ (0.1, 0.2, 0.4, or 0.8 Pa).

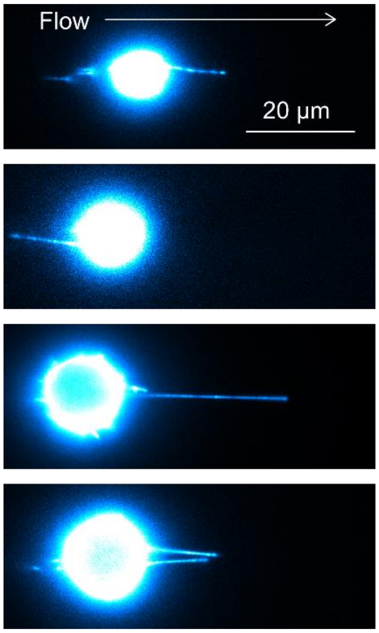


**Figure S10.** Formation of the tethers and slings on primary human CD34$^{pos}$-HSPCs rolling over E-selectin. Examples of the fluorescence images of CD44 (immunostained by Alexa-Fluor-647-conjugated anti-CD44 antibody, clone 515) captured during human CD34$^{pos}$-HSPC rolling over the surface-deposited rh E-selectin molecules. The cells were injected into the chambers at a shear stress of 2 dyne cm$^{-2}$ (0.2 Pa).

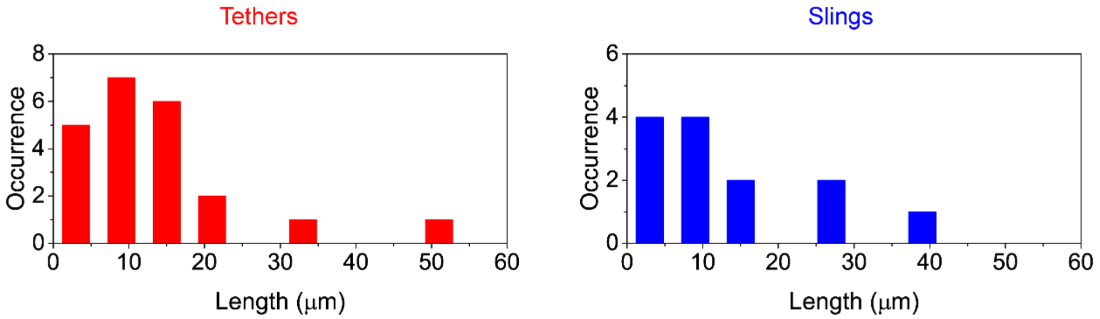


**Figure S11.** Length of the tethers and slings formed on primary human CD34$^{pos}$-HSPCs rolling over E-selectin. Frequency histograms of the length of tethers (red bars) and slings (blue bars) formed during the primary human CD34$^{pos}$-HSPCs rolling over the surface-deposited rh E-selectin. The cells were injected into the chambers at a shear stress of 2 dyne cm$^{-2}$ (0.2 Pa) as indicated.

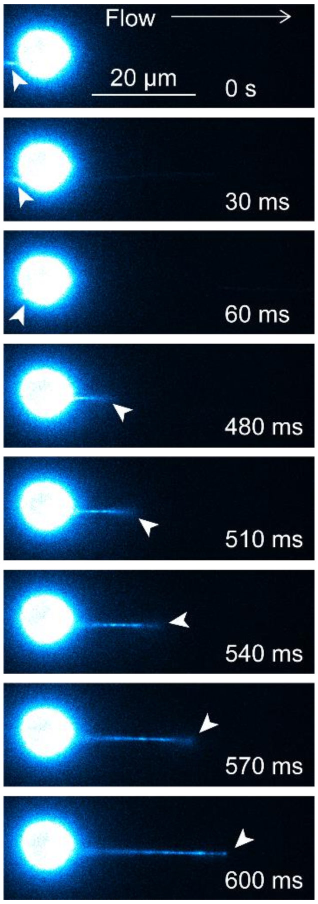


**Figure S12.** Conversion of a tether into sling observed for a primary human CD34$^{pos}$-HSPC rolling over E-selectin. Time-lapse fluorescence images of CD44 (immunostained by Alexa-Fluor-647-conjugated anti-CD44 antibody, clone 515) captured during cell rolling over the surface-deposited rh E-selectin molecules. The arrow heads show the tether that is converted into sling upon the detachment of the tethering point from the E-selectin surface. The cells were injected into the chambers at a shear stress of 2 dyne cm$^{-2}$ (Pa).

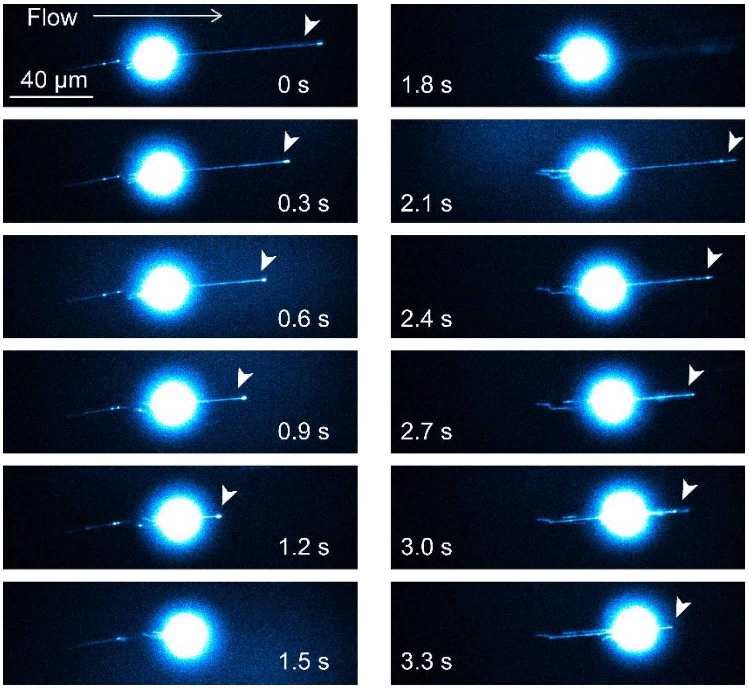


**Figure S13.** Retraction of the slings. Time-lapse fluorescence images of CD44 (immunostained by Alexa-Fluor-647-conjugated anti-CD44 antibody, clone 515) captured during cell rolling over the surface-deposited rh E-selectin molecules. The arrow heads show the slings with this retraction behavior. The cells were injected into the chambers at a shear stress of 8 dyne $cm^{-2}$ (0.8 Pa).

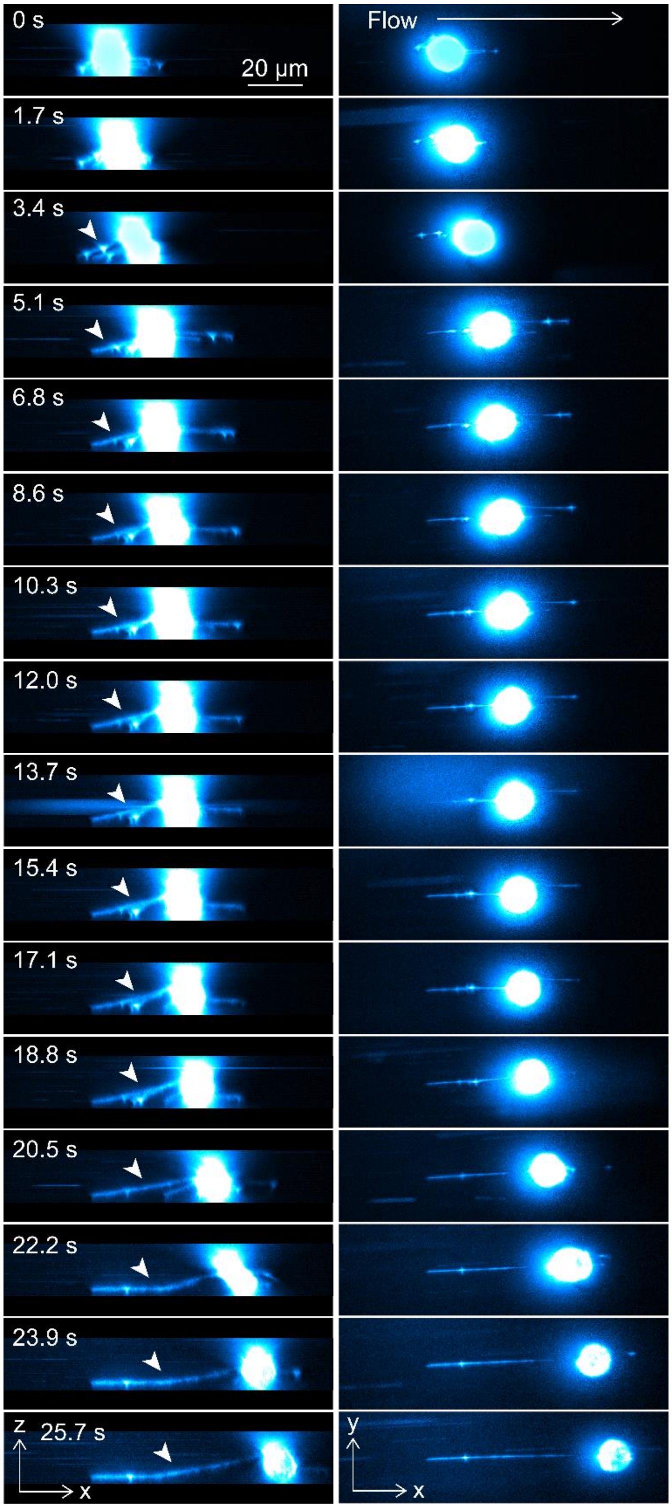


**Figure S14. 3D views of the elongation of the tether formed on a KG1a cell rolling over E-selectin.** Side view (left) and top view (right) of the 3D reconstructed time-lapse fluorescence images of CD44 (immunostained by Alexa-Fluor-488-conjugated anti-CD44 antibody, clone 515) captured during cell rolling over surface-deposited rh E-selectin molecules. The 3D images were reconstructed by recording fluorescence images of the cell at 57 different Z-axis positions with 0.5 μm step size. The arrowheads indicate the tether that shows elongation behavior. The cells were injected into the chambers at a shear stress of 2 dyne $cm^{-2}$ (0.2 Pa).

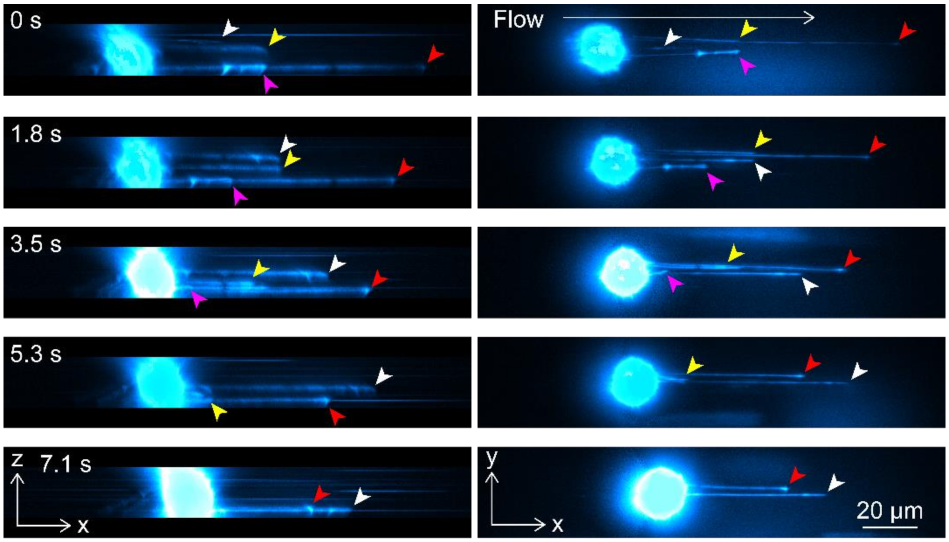


**Figure S15.** 3D views of the slings formed on a KG1a cell that show changes in the Z-axis positions during cell rolling over E-selectin. Side view (left) and top view (right) of the 3D reconstructed time-lapse fluorescence images of CD44 (immunostained by Alexa-Fluor-488-conjugated anti-CD44 antibody, clone 515) captured during cell rolling over the surface-deposited rh E-selectin molecules. The 3D images were reconstructed by recording fluorescence images of the cell at 59 different Z-axis positions with 0.5 μm step size. The arrow heads in white, yellow, red and magenta show four different slings formed during cell rolling. The cells were injected into the chambers at a shear stress of 2 dyne $cm^{-2}$ (0.2 Pa).

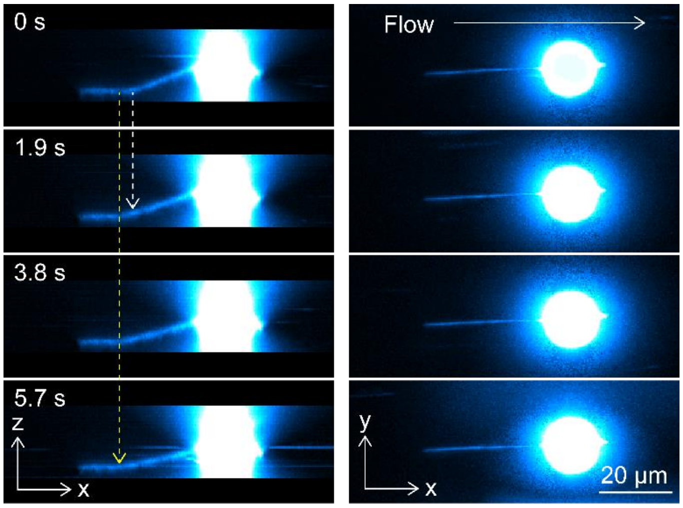


**Figure S16.** 3D views of the tether formed on a KG1a cell that show changes in the position of the anchoring points during cell rolling over E-selectin. Side view (left) and top view (right) of the 3D reconstructed time-lapse fluorescence images of CD44 (immunostained by Alexa-Fluor-488-conjugated anti-CD44 antibody, clone 515) captured during cell rolling over the surface-deposited rh E-selectin molecules. The 3D images were reconstructed by recording fluorescence images of the cell at 63 different Z-axis positions with 0.5 μm step size. The white and yellow arrows show the positions of the two anchoring points formed on the tether. The first anchoring point (white arrow) is detached during cell rolling, whereas the second anchoring point (yellow arrow) continues attaching to the surface E-selectin. The cells were injected into the chambers at a shear stress of 2 dyne $cm^{-2}$ (0.2 Pa).

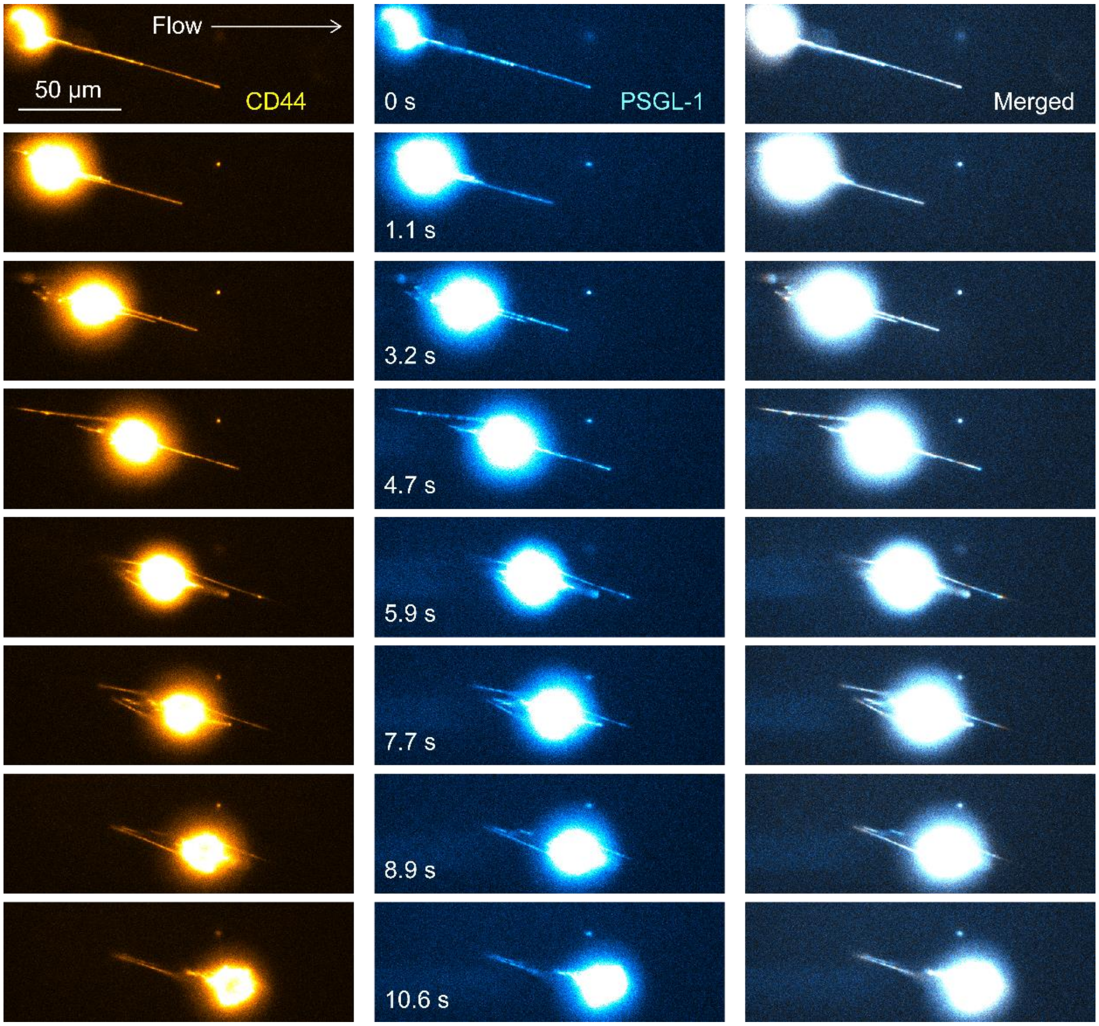


**Figure S17.** Colocalization of CD44 and PSGL-1 on the tethers and slings formed on a KG1a cell rolling over E-selectin. Time-lapse fluorescence images of CD44 (yellow, immunostained by Alexa-Fluor-647-conjugated anti-CD44 antibody, clone 515) and PSGL-1 (cyan, immunostained by Alexa-Fluor-488-conjugated anti-PSGL-1 antibody, clone KPL-1) captured during cell rolling over the surface-deposited rh E-selectin molecules. The merged images are displayed in the right panels. The cells were injected into the chambers at a shear stress of 2 dyne $cm^{-2}$ (0.2 Pa).

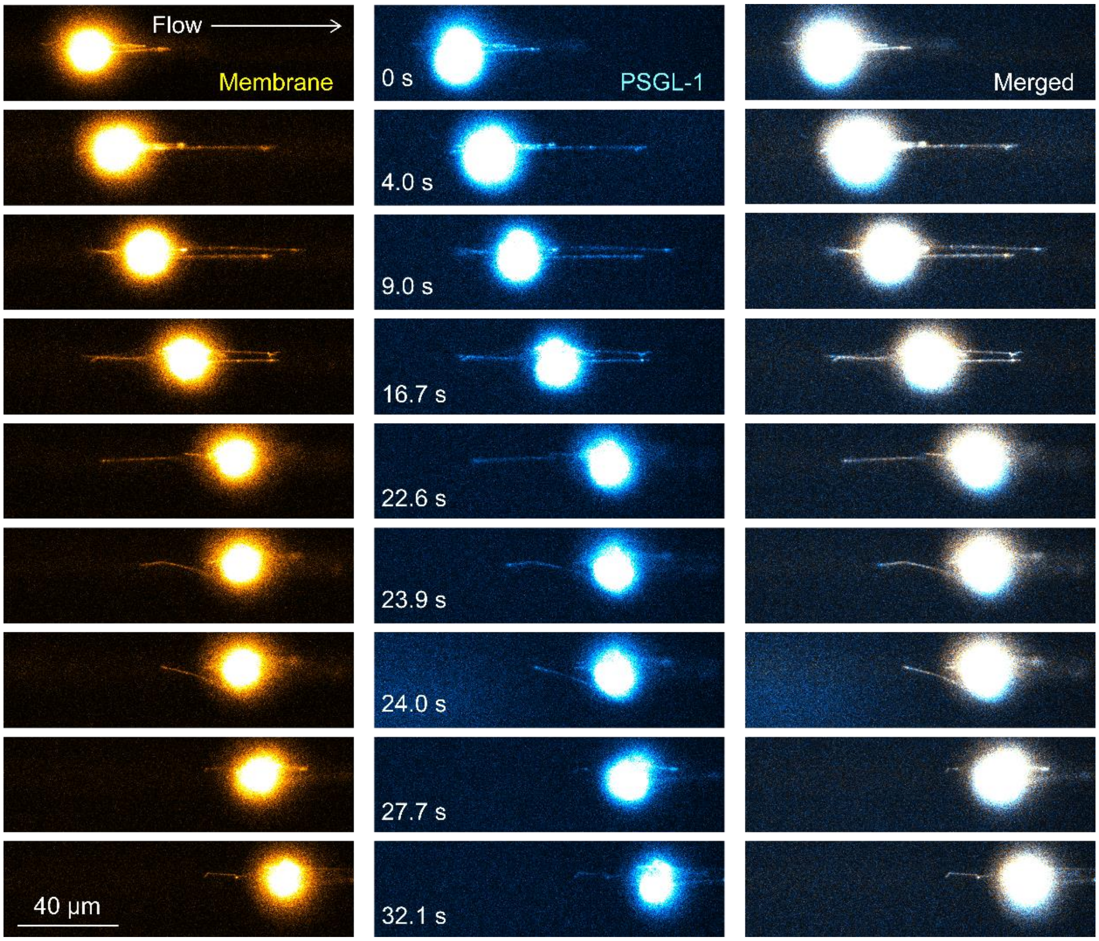


**Figure S18.** Colocalization of cell membrane and PSGL-1 on the tethers and slings formed on a KG1a cell rolling over E-selectin. Time-lapse fluorescence images of the cell membrane (yellow, stained by Vybrant DiO dye) and PSGL-1 (cyan, immunostained by Alexa-Fluor-647-conjugated anti-PSGL-1 antibody, clone KPL-1) captured during cell rolling over the surface-deposited rh E-selectin molecules. The merged images are displayed in the right panels. The cells were injected into the chambers at a shear stress of 2 dyne cm$^{-2}$ (0.2 Pa).

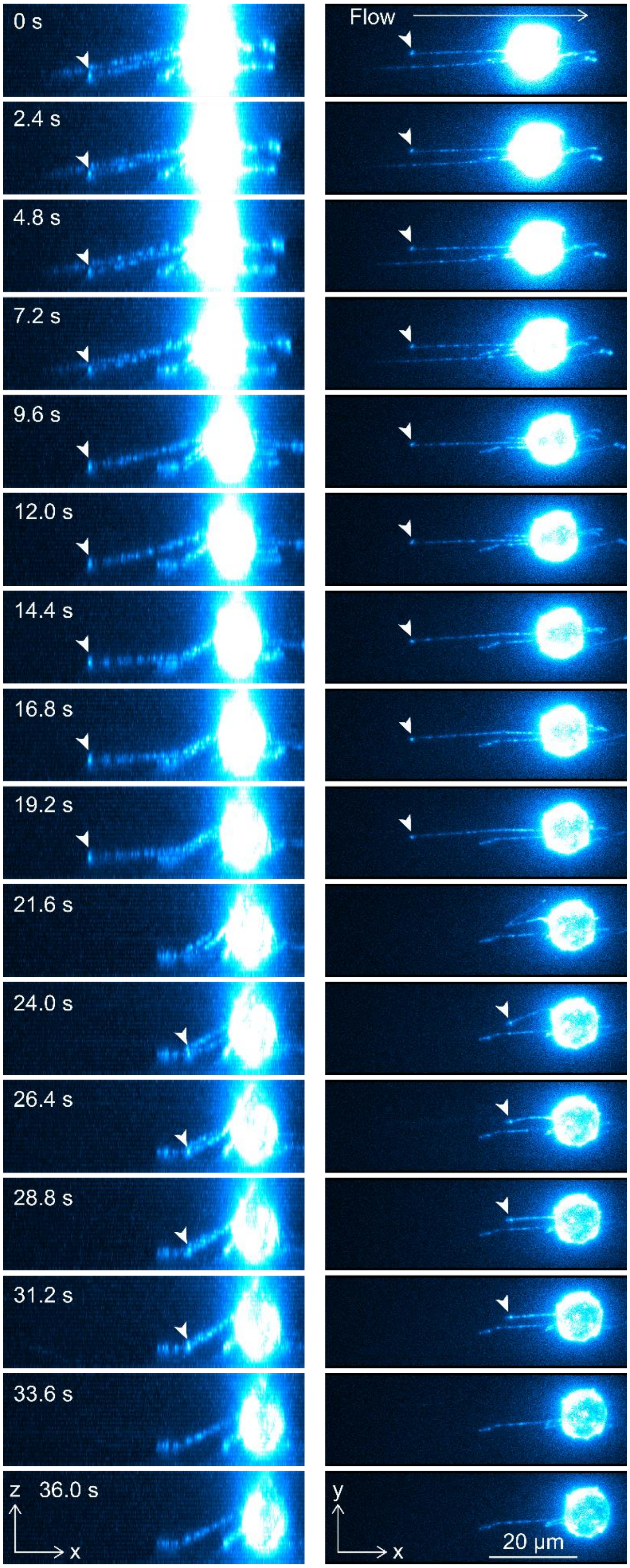
0 s
2.4 s
4.8 s
7.2 s
9.6 s
12.0 s
14.4 s
16.8 s
19.2 s
21.6 s
24.0 s
26.4 s
28.8 s
31.2 s
33.6 s
36.0 s
z
x
Flow
y
x
20 µm

**Figure S19.** 3D views of the tethers and slings formed on a KG1a cell that demonstrates clustering behavior of PSGL-1 at the tethering points during cell rolling over E-selectin. Side view (left) and top view (right) of the 3D reconstructed time-lapse fluorescence images of PSGL-1 (immunostained by Alexa-Fluor-555-conjugated anti-PSGL-1 antibody, clone KPL-1) captured during cell rolling over the surface-deposited rh E-selectin molecules. The 3D images were reconstructed by recording fluorescence images of the cell at 40 different Z-axis positions with 1.0 μm step size. The arrowheads show the bright fluorescence spots of PSGL-1 (i.e. clusters of PSGL-1) found at the tethering points. The cells were injected into the chambers at a shear stress of 2 dyne $cm^{-2}$ (0.2 Pa).

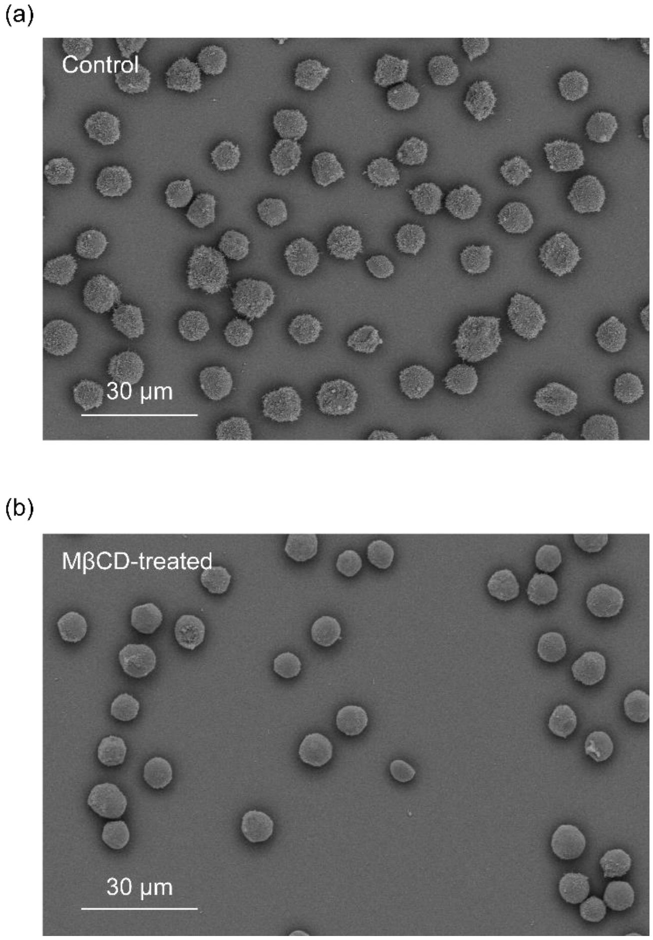


**Figure S20.** Scanning electron microscopy (SEM) images of KG1a cells. SEM images of (a) control and (b) methyl-β-cyclodextrin (MβCD)-treated KG1a cells. These are representative images of n=2 independent experiments.

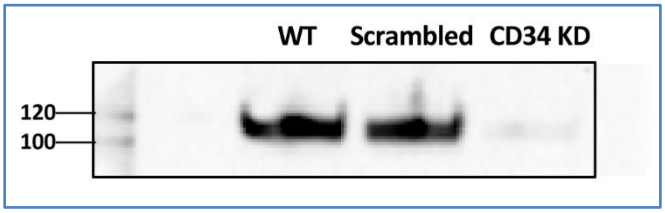


**Figure S21.** CD34 knockdown in KG1a cells. KG1a cells were transfected with either a scrambled control siRNA (Silencer Select, Life technologies; 4390843) or with siRNA specific for CD34 (Select, Life technologies; 4392420-s2644). Cells were collected after 48 h of transfection and subjected to Western blot analysis for CD34 protein expression. Lane 1: untreated KG1a cells; Lane 2: scrambled negative control siRNA; Lane 3: CD34 KD KG1a cells. Membrane bands were normalized using total cells protein. This is representative of n=5 independent experiments.

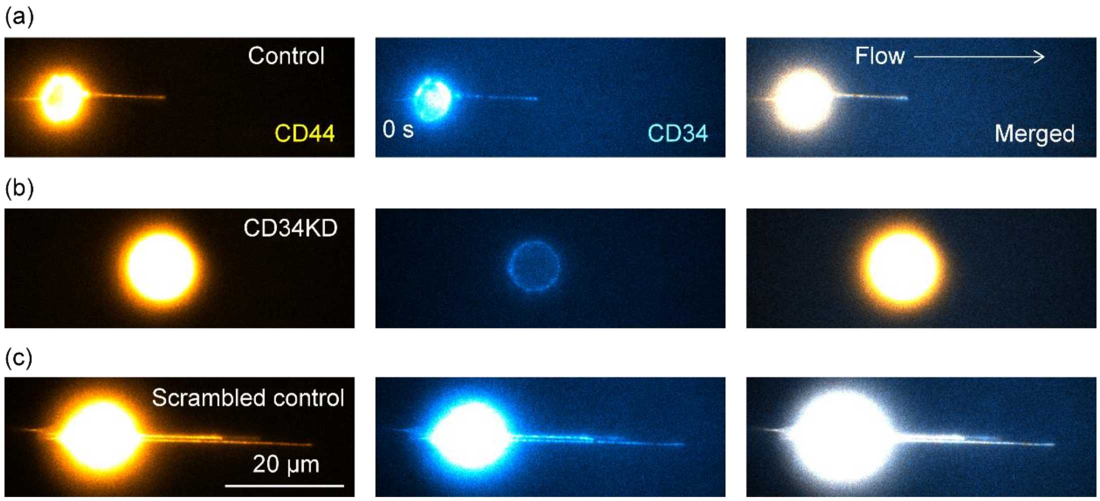


**Figure S22.** Effect of the knockdown of CD34 on the formation of tethers and slings on KG1a cells while rolling over E-selectin. Examples of the two-color fluorescence images of CD34 (cyan, immunostained by Alexa-Fluor-488-conjugated anti-CD34 antibody clone QBEND/10) and CD44 (yellow, immunostained by Alexa-Fluor-647-conjugated anti-CD44 antibody clone 515) on (a) control KG1a cell, (b) KG1a cell transfected with siRNA specific for CD34, and (c) KG1a cell transfected with scrambled control siRNA captured during cell rolling over the surface-deposited rh E-selectin molecules. The cells were injected into the chambers at a shear stress of 2 dyne cm$^{-2}$ (0.2 Pa). These images were captured using identical imaging conditions and were shown in an identical image contrast.

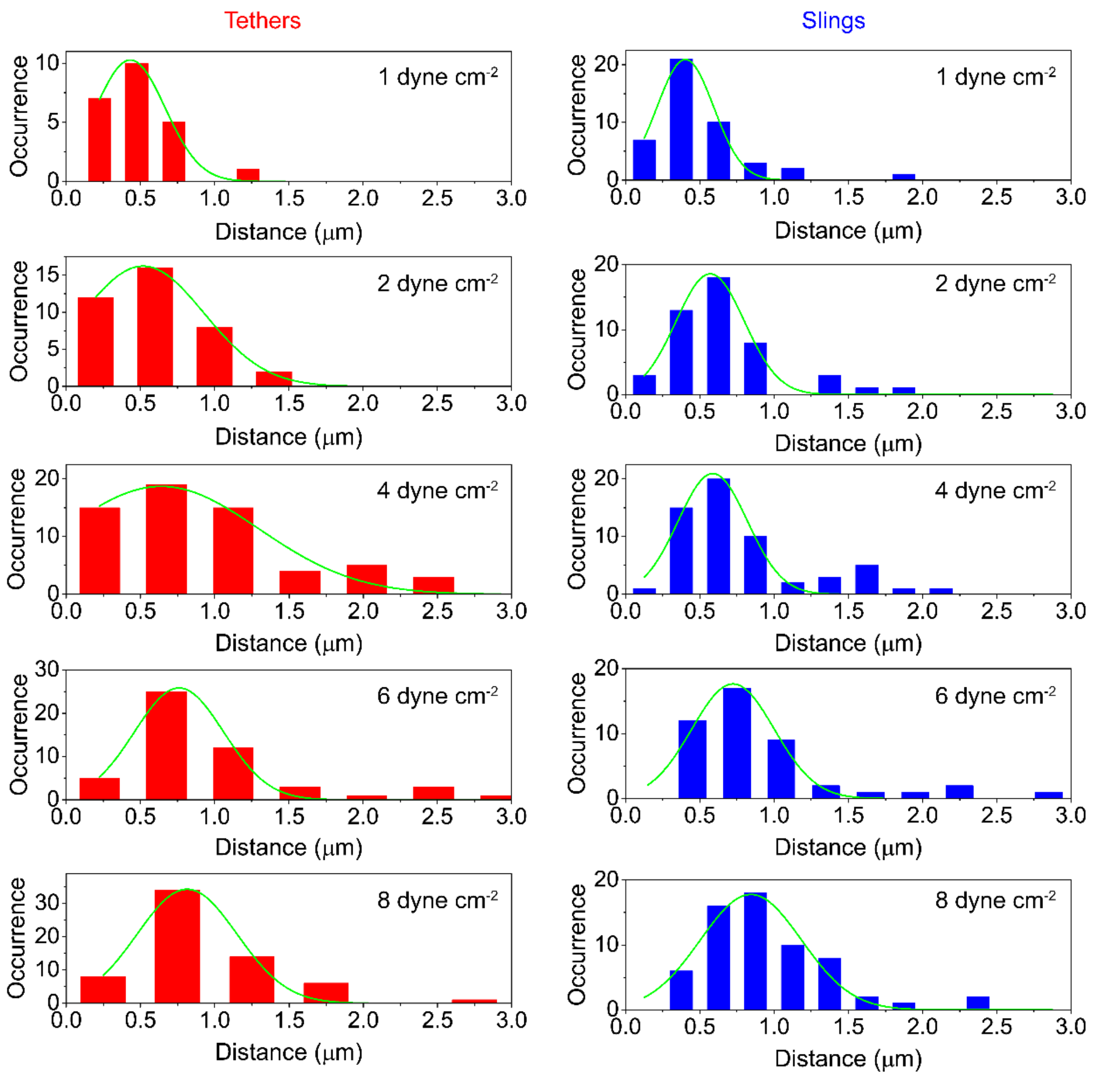


**Figure S23.** Distance between the PSGL-1 spots on the tethers and slings formed on KG1a cells rolling over E-selectin. Frequency histograms of the distance between the adjacent fluorescent spots of PSGL-1 (immunostained by Alexa-Fluor-555-conjugated anti-PSGL-1 antibody, clone KPL-1) on the tethers (red bars) and slings (blue bars) formed during the KG1a cells rolling over rh E-selectin. The cells were injected into the chambers at a shear stress of either 1, 2, 4, or 8 dyne cm$^{-2}$ (0.1, 0.2, 0.4, or 0.8 Pa).

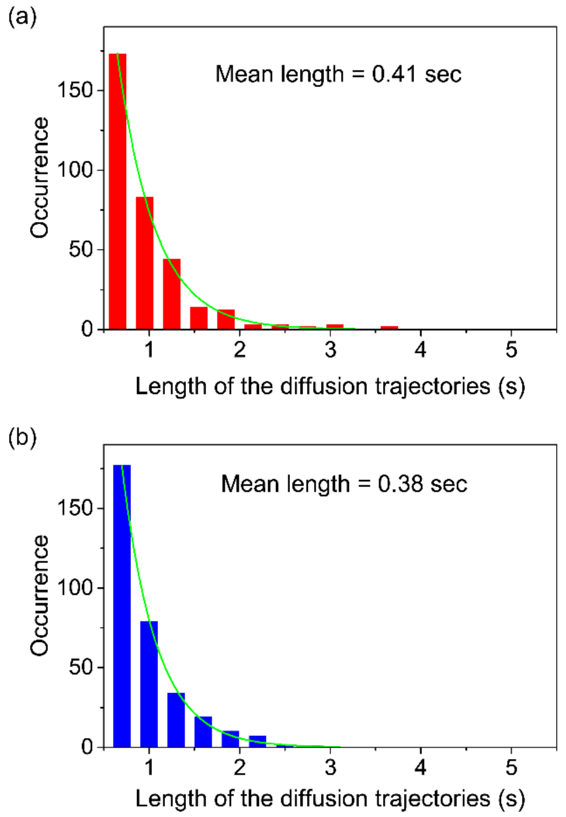


**Figure S24.** Length of the single-molecule diffusion trajectories of the PSGL-1 molecules on the tethers and slings formed on KG1a cells rolling over E-selectin. Frequency histograms of the length of the single-molecule diffusion trajectories of PSGL-1 molecules (immunostained by Alexa-Fluor-555-conjugated anti-PSGL-1 antibody, clone KPL-1) that show diffusional motion on the (a) tethers and (b) slings formed during the KG1a cells rolling over rh E-selectin. The solid lines show the fitting to single-exponential decaying function. The mean trajectory lengths (0.41s and 0.38 s for the trajectories obtained from the tethers and slings, respectively) were calculated using the decay constants obtained by the fits. The cells were injected into the chambers at a shear stress of 2 dyne $cm^{-2}$ (0.2 Pa).

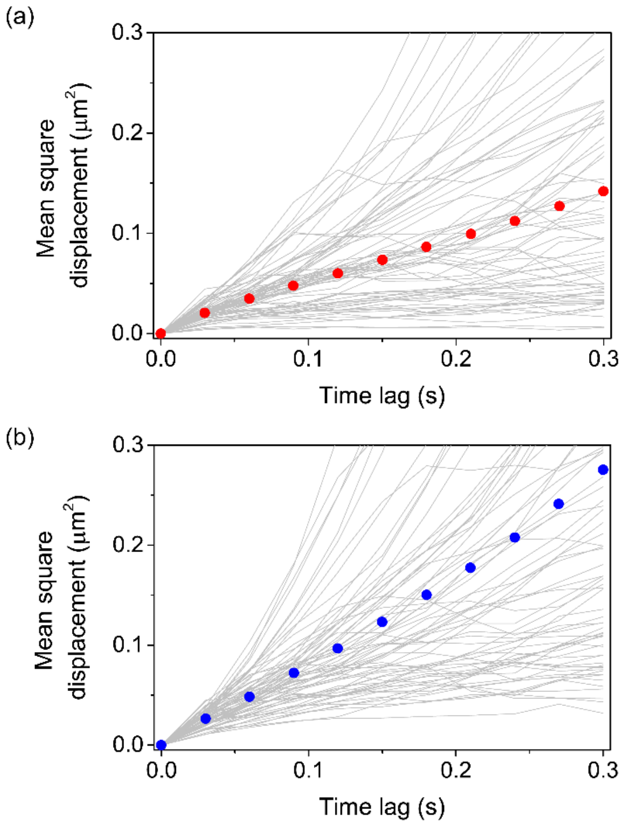


**Figure S25.** MSD vs time-lag plots obtained from the single-molecule diffusion trajectories of PSGL-1 on the tethers and slings. MSD vs time-lag plots obtained for PSGL-1 molecules (immunostained by Alexa-Fluor-555-conjugated anti-PSGL-1 antibody, clone KPL-1) that show diffusional motion on the (a) tethers and (b) slings formed during KG1a cells rolling over rh E-selectin. The grey lines show individual MSD vs time-lag plots obtained from individual single-molecule diffusion trajectories. The red and blue dots show mean MSD vs time-lag plots for the tethers and slings, respectively. The cells were injected into the chambers at a shear stress of 2 dyne cm$^{-2}$ (0.2 Pa).

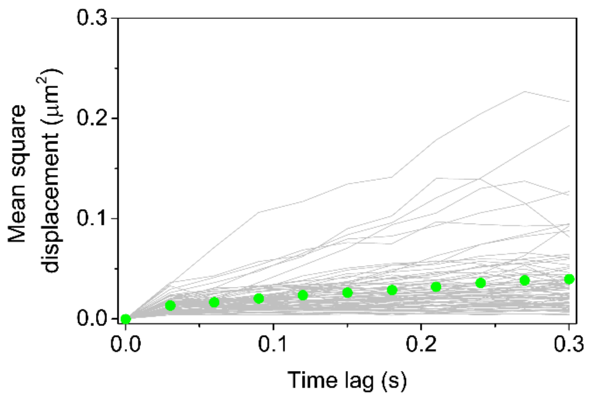


**Figure S26.** MSD vs time-lag plots obtained from the single-molecule diffusion trajectories of PSGL-1 localised on microvilli. MSD vs time-lag plots obtained for PSGL-1 molecules (immunostained by Alexa-Fluor-488-conjugated anti-PSGL-1 antibody, clone KPL-1) that are localized on microvilli in the control KG1a cells. The grey lines show individual MSD vs time-lag plots obtained from individual single-molecule diffusion trajectories. The green dots show mean MSD vs time-lag plot. The cells were injected into uncoated microfluidic chambers without any recombinant E-selectin molecules. The fluorescence images were captured in the absence of any external shear stress.

**Supporting Notes**

**1. Bivalency effect of the antibodies**

Antibodies have two binding sites to their epitopes. This may cause artificial clustering of the labeled molecules. Since we observed the discrete spatial distribution of the PSGL-1 molecule on the tethers and slings of the KG1a cell that are formed during the cell rolling over E-selectin, we investigated if this spatial distribution is a result of the artificial clustering of the immunostained PSGL-1 molecules due to the bivalency effect of the anti-PSGL-1 antibody. To that end, we immunostained PSGL-1 on KG1a cells using the Alexa-Fluor-555-conjugated Fab fragment of the anti-PSGL-1 antibody that has only one binding site to its epitope. The immunofluorescence image of PSGL-1 on the tethers and slings of the KG1a cells obtained using the Alexa-Fluor-555-conjugated Fab fragment of the anti-PSGL-1 antibody showed a spatial distribution of the PSGL-1 molecules on the tethers and slings very similar to that obtained using the Alexa-Fluor-555-conjugated anti-PSGL-1 antibody (i.e. whole antibody with two binding sites) (Figure SN1). This result demonstrates that the discrete spatial distribution of the PSGL-1 molecules on the tethers and slings are not the result of artificial clustering of PSGL-1 due to the bivalency effect of the antibody.

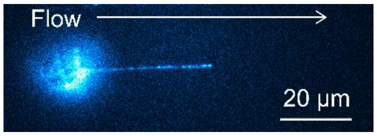


**Figure SN1.** Immunofluorescence image of PSGL-1 on KG1a cell captured using the Fab fragment of the anti-PSGL-1 antibody. The PSGL-1 molecules on the KG1a cells were immunostained by the Alexa-Fluor-555-conjugated Fab fragment of the anti-PSGL-1 antibody

(KPL-1 clone). The fluorescence image was recorded during the KG1a cells rolling over E-selectin at a shear stress of 2 dyne $cm^{-2}$ (0.2 Pa).

### 2. Density of CD44 on the tethers and slings

We calculated the number of the CD44 molecules per unit length of slings in a way similar to that for PSGL-1 (see Methods). The calculated densities of CD44 on the tethers and slings are in the range of 24 – 111 molecules $\mu m^{-1}$ (i.e. the distances between adjacent CD44 molecules on the tethers and slings are in the range of 9 – 42 nm). This result demonstrates quantitatively that the densities of CD44 on the tethers and slings are much higher than those of PSGL-1 (0.4 – 0.8 μm distances between adjacent molecules on the tethers and slings, i.e. 1.25 – 2.5 molecules $\mu m^{-1}$). The distances between adjacent CD44 molecules on the tethers and slings estimated by the density calculation (9 – 42 nm) are much shorter than the spatial resolution of our fluorescence microscope (approximately 200 nm according to Rayleigh criterion). Therefore, individual CD44 molecules on the tethers and slings cannot be resolved in our fluorescence imaging experiment. This is consistent with the contiguous distribution of CD44 observed in our imaging experiment. The density of CD44 molecules on the tethers and slings decreased with an increase in the shear stress (Figure SN2). Similar shear stress-dependent density was observed for PSGL-1 (Figure 6g).

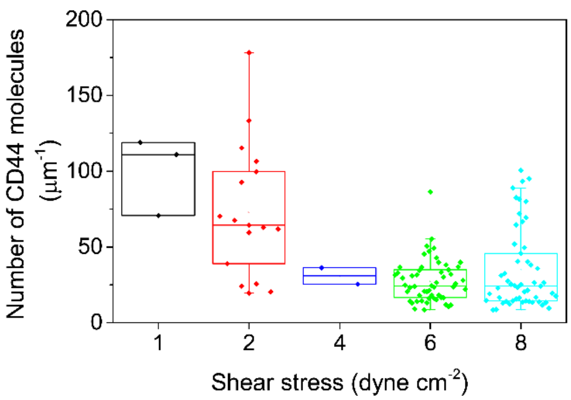


**Figure SN2.** Number of CD44 molecules per unit length of slings. The CD44 molecules on the KG1a cells were immunostained by the Alexa-Fluor-647-conjugated anti-CD44 antibody (515 clone). The fluorescence images were recorded during KG1a cell rolling over E-selectin at shear stresses of 1, 2, 4, 6, and 8 dynes cm$^{-2}$ (0.1, 0.2, 0.4, 0.6, and 0.8 Pa).

#### 3. Comparison of E-selectin and P-selectin in terms of the tether formation

We observed the formation of the tethers with mean lengths of 18 – 30 µm at shear stresses ranging between 1 and 8 dyne cm$^{-2}$ (0.1 and 0.8 Pa) (Figure 3b, 6f). Although the formation of tethers has been reported for neutrophils rolling over P-selectin,[1] relatively short tethers (mean lengths of approximately 10 µm) were observed only at high shear stresses (above 6 dyne cm$^{-2}$ (0.6 Pa)).[2] Although we cannot rule out the possibility that cell type dependent elastic properties of cell membranes may affect the tethering behaviour, it is more likely that the difference is due to the strength of selectin-ligand interactions. Our observations suggest that E-selectin-ligand interactions are much stronger than P-selectin-ligand interactions. Since rapture force of E-selectin-ligand interactions observed for polymorphonuclear leukocytes is similar to that of P-selectin-ligand interactions,[3] it is likely that the difference in the binding strength between E-selectin and P-selectin is a result of different dissociation constants against their ligands. The stronger binding (i.e. smaller dissociation rate) of the tethers resists shear stresses for longer periods of time before they dissociate from the surface selectins, and thus causes the formation of longer tethers. This stronger binding is consistent with previous studies from our lab illustrating that the $k_{off}$ of binding between selectin ligands and E-selectin is low.[4, 5]

**4. Elastic property of the tethers and slings and its contribution to cell rolling**

The very bright and contiguous fluorescence of CD44 from the tethers and slings (Figure 3c) allows detailed discussion about the dynamic behaviour of the tethers and slings. Our live-cell fluorescence imaging experiment clearly captured the flipping of the broken tethers from the rear side of the KG1a cell to its front side during rolling on E-selectin (Figure 3c). The images demonstrate that the flipping of the tethers occurred at a time scale of approximately 240 ms. This time scale is longer than that reported for neutrophils rolling on P-selectin (less than 100 ms),[2] although similar tethers and slings were observed in both cell types. Given much larger shear stress (10 dyne $cm^{-2}$ (1 Pa)) used in the experiment on neutrophils compared with our experiment (Figure 3c, 6 dyne $cm^{-2}$ (0.6 Pa)), the observed difference is likely to be due to the different flow velocities in these experiments rather than different elastic properties of the tethers formed in these experiments. These results indicate that the elastic properties of the tethers and slings that make these structures so flexible are similar in different cell types, implying the existence of common mechanisms in the formation of tethers and slings.

**5. Colocalisation of CD44 and membrane tethers/slings on MβCD-treated KG1a cells**

The two-color fluorescence imaging experiment of the membrane stain and CD44 on the control KG1a cells showed perfect spatial colocalization of CD44 and the cell membrane in the tethers and slings (Figure 2a), and thus the spatiotemporal behavior of the tethers and slings (e.g. length of the tethers and slings at different applied shear stresses) can be characterized by analyzing fluorescence images obtained for CD44. The MβCD treatment of the cells may alter the spatial localization of CD44, which may affect the analysis of the dynamic behavior of the tethers and slings. We evaluated this by capturing two-color fluorescence images of the membrane stain and CD44 on the MβCD-treated KG1a cells (Figure SN3). While the tethers and slings of the MβCD-treated cells formed during cell rolling over E-selectin were much shorter than those of the control cells (Figure 5e), we found perfect spatial colocalization of the membrane stain and CD44 (Figure SN3). This result confirms that the spatiotemporal behavior of the tethers and slings on the MβCD-treated cells can be characterized by analyzing fluorescence images obtained for CD44.

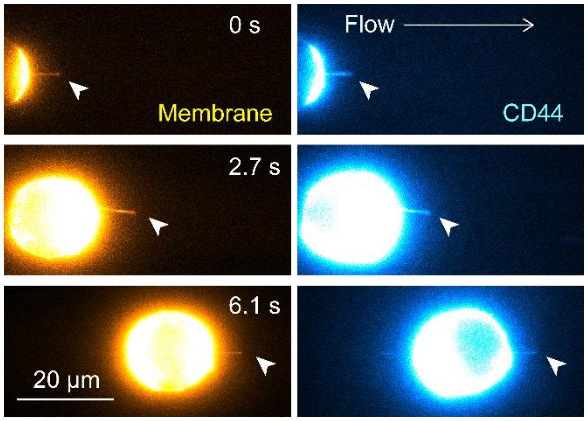


**Figure SN3. Spatial colocalization of CD44 and membrane tethers/slings on the MβCD-treated KG1a cells.** Two-color fluorescence images of the cell membrane (stained by Vybrant DiO dye) and CD44 (immunostained by Alexa-Fluor-647-conjugated anti-CD44 antibody clone 515) captured during the MβCD-treated KG1a cell rolling over the surface-deposited rh E-selectin molecules. White arrowheads show slings formed during the cell rolling. The cells were injected into the chambers at a shear stress of 2 dyne $cm^{-2}$ (0.2 Pa).

### 6. Effect of the retraction of the slings and rolling of the cells on the single-molecule diffusion analysis of PSGL-1

The single-molecule tracking analysis of the PSGL-1 molecules on the slings could be affected by the motion of the slings (i.e. either stable or retracting) and the cell (i.e. either rolling within the acquisition time of the diffusion trajectory or not). Thus, we split the diffusion trajectories into three categories and calculated the mean square displacement (MSD) versus time lag plots separately.

A. PSGL-1 molecules diffusing on the stable slings without the effect of the cell rolling.

B. PSGL-1 molecules diffusing on the retracting slings without the effect of the cell rolling.

C. PSGL-1 molecules diffusing on the stable slings with the effect of the cell rolling.

While we observed slight differences between the three cases, in principle, all the MSD versus time lag plots showed linear relationship (Figure SN4). This result confirms the random-mode diffusion of the PSGL-1 molecules on the slings. The effect of the retraction of the slings and the cell rolling is negligible in our MSD analysis probably because of the decoupling of the time scale of the diffusional motion of the PSGL-1 molecules and the motion of the slings and the cells.

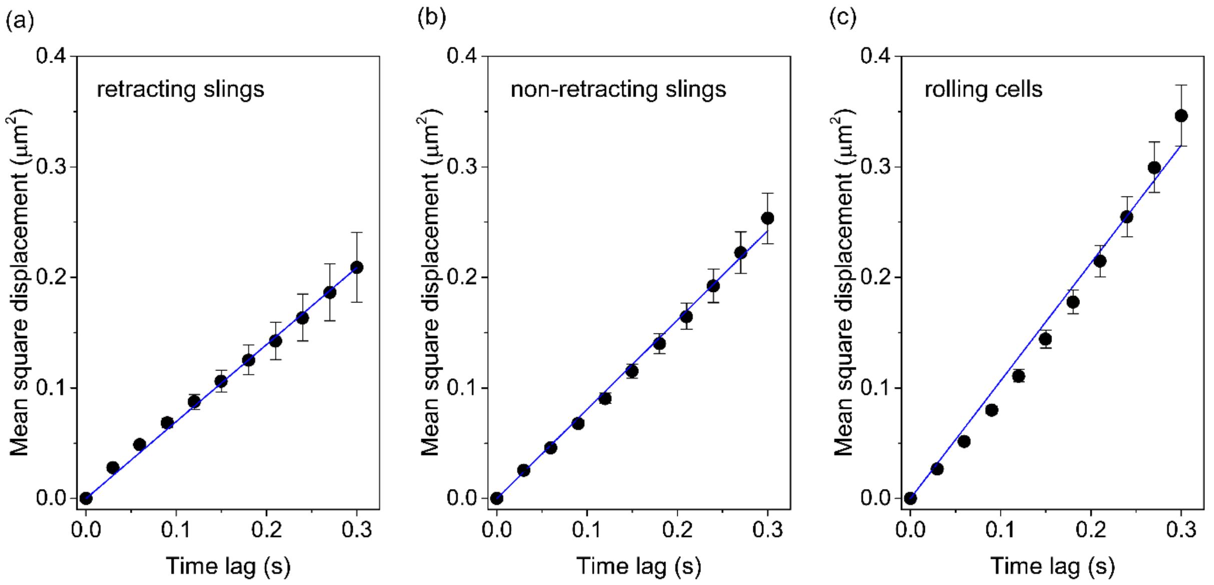


**Figure SN4. MSD analysis of the PSGL-1 molecule diffusing on the slings.** The MSD versus time lag plots were calculated using single-molecule diffusion trajectories of the PSGL-1 molecules diffusing on the (a) retracting slings without the effect of the cell rolling, (b) stable slings without the effect of the cell rolling and (c) stable slings with the effect of the cell rolling. The error bars show the standard errors of the mean determined by 47, 169 and 104 MSD plots obtained for the PSGL-1 molecules diffusing on the retracting slings, non-retracting slings and stable slings with the effect of the cell rolling, respectively.

### 7. Effect of the surface on the single-molecule tracking analysis and the interpretation of the obtained data

The single-molecule imaging experiment of the PSGL-1 molecules that are localized on the microvilli of the control KG1a cells (i.e. not rolling cells) was conducted by placing the immunostained cells on the surface of the microfluidic chamber. Although we did not deposit any molecules that have a specific interaction with the PSGL-1 molecules (e.g. E-selectin), the PSGL-

1 molecule may interact with the surface in a nonspecific manner, and therefore the diffusional motion of the PSGL-1 may be affected. We investigated this effect by analysing the single-molecule diffusion trajectories obtained from both the bottom and top surfaces of the cells. To minimize the effect of the refractive index mismatch between the cell samples and the immersion media, we used the silicone immersion objective lens (60× NA = 1.3, UPLSAPO60XS2) for this imaging experiment. The MSD versus time lag plots obtained from these experiments showed very similar behavior, including smaller diffusion coefficient compared with that obtained from the PSGL-1 molecules on the tethers and slings and confined-mode diffusion (Figure SN5a). Quantitative analysis of the MSD versus time lag plots revealed that the PSGL-1 molecules localized on the microvilli at both the bottom and top surface of the cells show similar diffusion coefficient and the confined size (Figure SN5b, c). These results confirm that there is a negligible effect of the surface on the diffusional motion of the PSGL-1 molecules localised on microvilli. We used the data obtained from the bottom surface of the cells for the analysis as we obtained slightly better quality of the single-molecule fluorescence images of the PSGL-1 molecules from the bottom surface of the cells.

The diffusion coefficient of the PSGL-1 molecules localized on the microvilli of the control KG1a cells obtained from the MSD analysis (0.1 $\mu m^2\ s^{-1}$) is approximately 30 fold larger than the diffusion coefficient of PSGL-1 reported previously using fluorescence recovery after photobleaching (FRAP) technique (0.003 $\mu m^2\ s^{-1}$).[6] Given the different time and length scales of the diffusional motion captured by these two methods, it is very likely that we captured the motion of microvilli rather than the diffusional motion of the PSGL-1 molecules in our single-molecule fluorescence imaging experiment (i.e. the PSGL-1 molecules stay at the tip of microvilli during the image acquisition and therefore we capture the motion of the microvilli through the fluorescence signal of the PSGL-1 molecules). This is strongly supported by the fact that the size of the confinement area (0.29 μm$^2$ that corresponds to the confinement length of approximately 0.54 μm) estimated by the MSD analysis is close to the size of the area expected to be covered by the microvilli since our SEM experiments on the KG1a cells revealed that the length of the microvilli is in the range of several hundred of nanometers (Figure 5c). Also, the length of the microvilli on neutrophils has been estimated to be about 0.3 μm.[7] Therefore, the real diffusion coefficient of the PSGL-1 molecules localised on the microvilli would be much smaller than our estimation by the MSD analysis of the single-molecule diffusion trajectories and may be closer to the value determined by the FRAP technique. The difference in the diffusion coefficient obtained in our experiment and in the previous study may also be partly explained by differences from cell to cell.

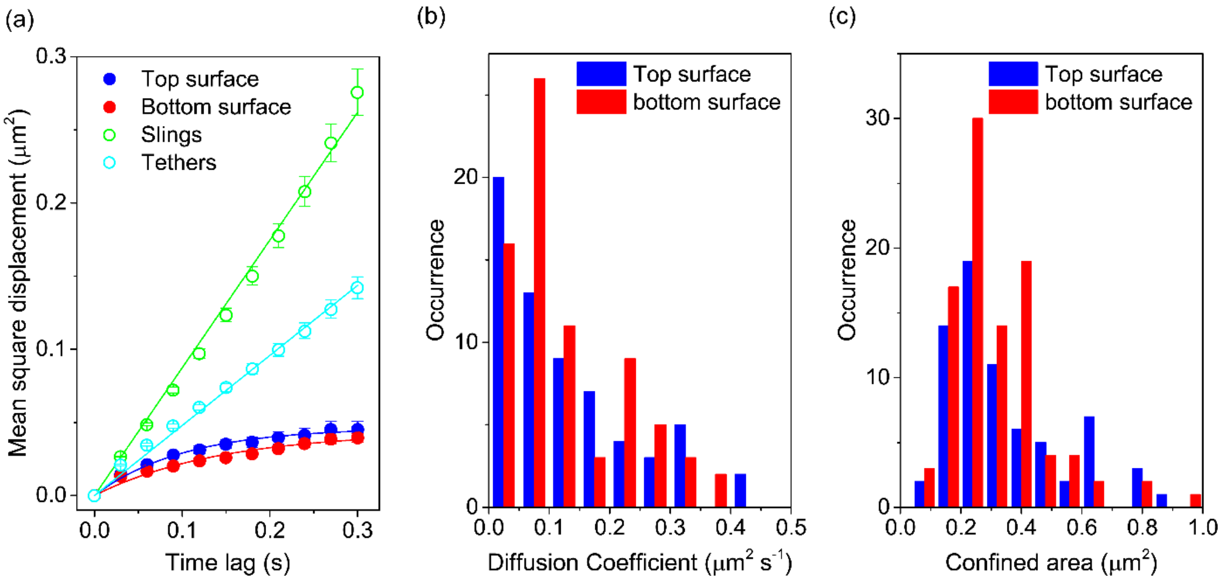


**Figure SN5. MSD analysis of the PSGL-1 molecule localised on the microvilli.** (a) MSD versus time lag plots calculated using single-molecule diffusion trajectories of the PSGL-1 molecules localized on the microvilli of the control KG1a cells at the top (blue) and bottom (red) surface of the cells. The error bars show the standard errors of the mean determined by 126 and 129 MSD plots obtained for the PSGL-1 molecules localized on the microvilli of the control KG1a cells at the top and bottom surface of the cells. The MSD versus time lag plots obtained from the single-molecule diffusion trajectories of the PSGL-1 molecules diffusing on the tethers (cyan) and slings (green) are also displayed as a comparison. Frequency histograms of the (b) diffusion coefficient and (c) confined area of the PSGL-1 molecules on the microvilli of the control KG1a cells at the top (blue) and bottom (red) surface of the cells.

**Captions for Movies S1 to S7**

**Movie S1:** Time-lapse fluorescence images of CD44 on a KG1a cell perfused into the rh E-selectin-deposited microfluidic chamber at a shear stress of 1 – 8 dyne $cm^{-2}$ (1 – 8 Pa). The CD44 molecules were immunostained by Alexa-Fluor-647-conjugated anti-CD44 antibody, clone 515. Scale bar = 20 μm.

**Movie S2:** Time-lapse fluorescence images of CD44 on a KG1a cell perfused into the rh E-selectin-deposited microfluidic chamber that shows the conversion of a tether into sling. The CD44 molecules were immunostained by Alexa-Fluor-647-conjugated anti-CD44 antibody, clone 515. The cells were injected into the chambers at a shear stress of 6 dyne $cm^{-2}$ (0.6 Pa). Scale bar = 20 μm.

**Movie S3:** Time-lapse fluorescence images of CD44 on a KG1a cell perfused into the rh E-selectin-deposited microfluidic chamber that shows the retraction of the sling. The CD44 molecules were immunostained by Alexa-Fluor-647-conjugated anti-CD44 antibody, clone 515. The cells were injected into the chambers at a shear stress of 8 dyne $cm^{-2}$ (0.8 Pa). Scale bar = 20 μm.

**Movie S4:** Time-lapse fluorescence images of PSGL-1 on a KG1a cell perfused into the rh E-selectin-deposited microfluidic chamber that shows the discrete spatial distribution of the PSGL-1 molecules on the tether. The PSGL- molecules were immunostained by Alexa-Fluor-555-conjugated anti-PSGL-1 antibody, clone KPL-1. The cells were injected into the chambers at the shear stress of 2 dyne $cm^{-2}$ (0.2 Pa). Scale bar = 10 μm.

**Movie S5:** **T**ime-lapse fluorescence images of PSGL-1 on a KG1a cell perfused into the rh E-selectin-deposited microfluidic chamber that shows the discrete spatial distribution of the PSGL-1 molecules on the sling. The PSGL- molecules were immunostained by Alexa-Fluor-555-conjugated anti-PSGL-1 antibody, clone KPL-1. The cells were injected into the chambers at a shear stress of 2 dyne $cm^{-2}$ (0.2 Pa). Scale bar = 10 μm.

**Movie S6:** Time-lapse fluorescence images of CD44 on KG1a cells perfused into the rh E-selectin-deposited microfluidic chamber that shows the formation of the tethers and slings on all the rolling cells. The CD44 molecules were immunostained by Alexa-Fluor-647-conjugated anti-CD44 antibody, clone 515. The cells were injected into the chambers at a shear stress of 4 dyne $cm^{-2}$ (0.4 Pa). Scale bar = 20 μm.

**Movie S7:** Time-lapse fluorescence images of PSGL-1 on a KG1a cell perfused into the rh E-selectin-deposited microfluidic chamber that shows the merger of multiple tethers. The PSGL-molecules were immunostained by Alexa-Fluor-555-conjugated anti-PSGL-1 antibody, clone KPL-1. The cells were injected into the chambers at a shear stress of 2 dyne $cm^{-2}$ (0.2 Pa). Scale bar = 5 μm.
